# Pathogen and host adapt pH responses during enteric infection

**DOI:** 10.1101/2024.01.05.573998

**Authors:** Sarah E. Woodward, Laurel M.P. Neufeld, Jorge Peña-Díaz, Wenny Feng, Antonio Serapio-Palacios, Isabel Tarrant, B. Brett Finlay

**Affiliations:** Department of Microbiology and Immunology, University of British Columbia, Vancouver, BC, Canada; Michael Smith Laboratories, University of British Columbia, Vancouver, BC, Canada; Department of Biochemistry and Molecular Biology, University of British Columbia, Vancouver, BC, Canada

## Abstract

Enteric pathogens navigate distinct regional micro-environments within the intestine which cue important adaptive behaviours. We investigated the response of *Citrobacter rodentium*, a model of human pathogenic *Escherichia coli* infection, to regional gastrointestinal pH. We found that small intestinal pH (4.4-4.8) triggered virulence gene expression and altered cell morphology, supporting initial intestinal attachment, while higher pH, representative of *C. rodentium*’s replicative niches further along the intestine, supported pathogen growth. Gastric pH, a key barrier to intestinal colonization, caused significant accumulation of intra-bacterial reactive oxygen species, inhibiting growth of *C. rodentium* and related human pathogens. Within-host adaptation increased gastric acid survival, which may be due to a robust acid tolerance response induced at colonic pH. However, we also found that host gastric pH decreases post-infection, corresponding to increased serum gastrin levels and altered host expression of acid secretion-related genes. Similar responses following *Salmonella* infection may indicate a protective host response to limit further pathogen ingestion. Together, we highlight adaptive pH responses as an important component of host-pathogen co-evolution.

## Introduction

Gastrointestinal pH is a key component of gut physiology, known to vary across regions of the intestine (4,5). In humans, intestinal pH is characterized by a progressive increase from the duodenum to the distal ileum (pH 6.0-7.5), a drop at the cecum to pH 6.4, followed by increasing pH from the proximal to the distal colon (pH 7-7.5; Figure 1A) (4–6). In mice, reported intestinal pH is much lower, with the intestinal contents of female BALB/c mice ranging from 2.98 in the stomach to 5.24 in the ileum (Figure 1A) (7). In both species, maintaining intestinal pH is important to homeostasis and gut function. Disruption to intestinal ion transport is associated with spontaneous onset of colitis symptoms in mice, and increased susceptibility to induced colitis (8–10). pH also influences which microbial species colonize the gut, with mildly acidic pH resembling proximal colon conditions inhibiting the growth of *Bacteroides spp*. (11–13). Therefore, maintenance of gut pH is important to preventing overgrowth of colonizing bacterial, whether commensal or pathogenic (11,12).

**Figure 1.**
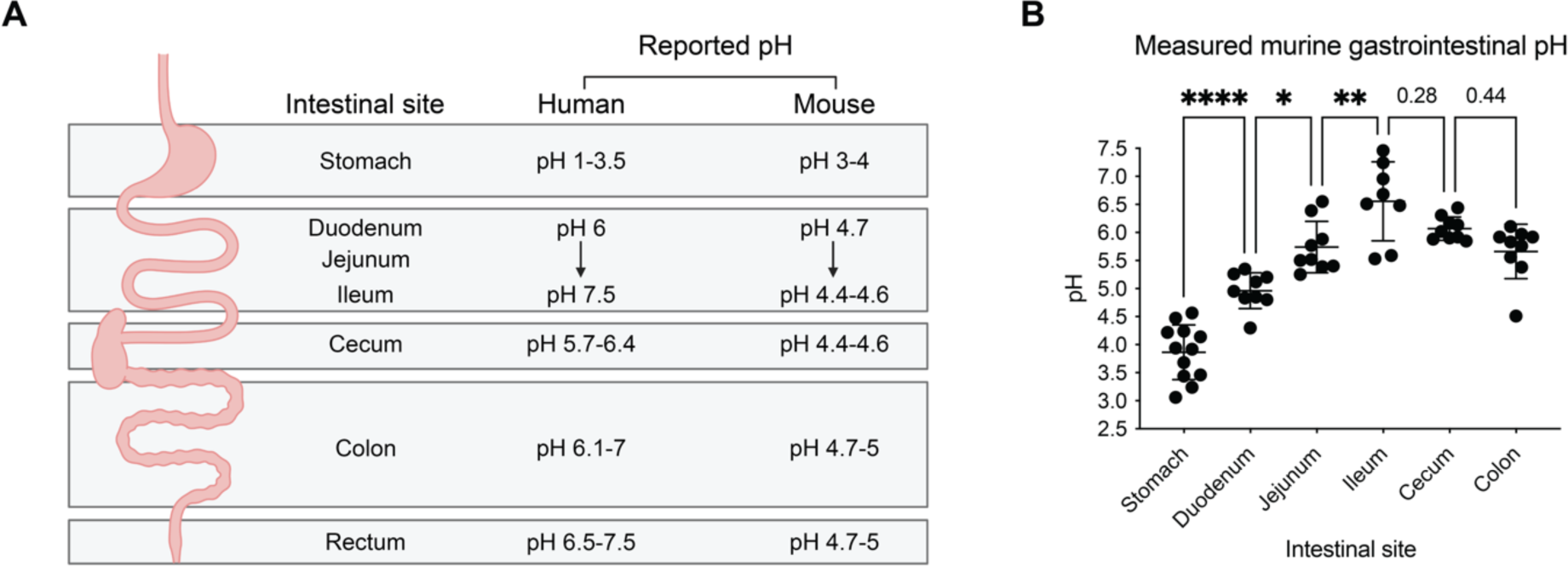
Gastrointestinal pH varies across gut regions. A) Average lumenal pH of gut regions in murine and human gastrointestinal tracts, as reported in the literature (4–6). B) Lumenal pH measured across the gut of uninfected C57BL/6 mice (N = 8-12). Error represents mean +/- SD. Statistical analysis represents a one-way ANOVA with Tukey’s multiple comparisons test.

Intestinal pH is often overlooked with regard to enteric infection, despite its potential role in affecting infection dynamics, host-pathogen interactions, and possibly even therapeutic success. By definition, diarrheal disease agents interact with the GI environment to establish infection. Pathogenic *Escherichia coli*, including Enteropathogenic *E. coli* (EPEC) and enterohemorrhagic *E. coli* (EHEC), are a diverse group of diarrheal disease agents which result in diarrhea, fever, vomiting, and even death in infected hosts (15,16). *Citrobacter rodentium*, a related pathogen that shares similar virulence factors, is used to model murine EPEC/EHEC infection (17–19). *Citrobacter rodentium* naturally infects mice, causing transmissible murine colonic hyperplasia (a form of colitis) with diarrheal disease symptoms (20). All three pathogens are transmitted through the fecal-oral route and use a type III secretion system (T3SS) for successful colonization (17,21–25), translocating protein effectors directly into host cells for bacterial control over host cell processes (26).

It is clear that EPEC/EHEC/*C. rodentium* respond to intestinal pH during host colonization. In particular, the extreme acidic conditions of the gastric environment are a major bottleneck to both *C. rodentium* and EHEC colonization (27–30). Furthermore, within the small intestine *C. rodentium* responds to alkaline bicarbonate ions secreted by the host to neutralize acidic chyme by upregulating genes involved in initial epithelial attachment (31,32). Despite this work, much remains unclear about how EPEC/EHEC/*C. rodentium* respond to regional differences in gastrointestinal pH. Furthermore, while the GI environment is known to change as a consequence of diarrheal disease, little investigation has been done to profile intestinal pH post-infection (1).

This study investigates the behaviour and virulence of *C. rodentium* in response to variations in host intestinal pH. We profiled pH across the intestine of one of the most commonly used laboratory mouse strains (C57BL/6) to define a range of physiologically relevant pH encountered by enteric pathogens upon initial host contact. We identified pH-specific regulation of the T3SS and epithelial attachment by *C. rodentium*, as well as significant changes to stress adaptation. Additionally, within-host adaptation increased *C. rodentium*’s acid tolerance, stimulated by colonic pH conditions. The pH of the gut also changed during established *C. rodentium* infection, with a significant drop in gastric pH which correlated with pathogen burden. This effect was consistent across additional models of intestinal damage, suggesting a possible host defense mechanism against further pathogen ingestion. Together, these data emphasize the interplay between pathogens and host intestinal pH in the context of enteric infection.

## Materials and Methods

### Bacterial strains and culture conditions

Bacterial strains are listed in Table S1. A fresh bacterial stock was used for all assays as follows: two days in advance bacteria were streaked onto a fresh LB agar plate from frozen (−70 °C in glycerol) and grown overnight at 37 °C. A single colony was re-inoculated into LB broth and grown overnight at 37 °C, shaking. To test the response of bacteria to physiological pH, LB (Lysogeny Broth, Sigma) or DMEM (Dulbecco’s Modified Eagle Medium, Hyclone) was adjusted to physiological pH by the addition of 37% HCl or 40% NaOH. Due to the high buffering capabilities of DMEM, media was pH-adjusted fresh for use within 12 hours. pHs tested were: 3.5, 4.0, 4.4, 4.8, 5.7, 6.0, 6.4, 7.0, and 7.5.

### Mouse infections and *in vivo* pH measurements

Animal experiments were performed in accordance with the guidelines of the Canadian Council on Animal Care and the University of British Columbia (UBC) Animal Care Committee according to Animal Care Protocols A20-0187 and A17-0228. C57BL/6 and BALB/c mice were ordered from Jackson Laboratory (Bar Harbor, ME) and maintained in a specific pathogen-free facility at UBC on a 12-hour light-dark cycle. Mice were allowed to acclimatize to the facility for one week following arrival. All pH measurements and infection models were run after the acclimatization period.

pH measurements were performed on fresh, undiluted intestinal content, using an Elite pH Spear Pocket Tester (Thermo Scientific). Measured pH represents the average of 2-3 readings per sample. All bacterial colonization experiments were done in 7-week-old female C57BL/6 mice fed the Picolab Rodent Diet 20 chow. Mice were monitored daily throughout the 4-8 day infections for weight loss and clinical symptoms.

*C. rodentium* infections were carried by gavaging mice orally with 10^8^ colony forming units (CFU) of *C. rodentium* DBS100 from overnight culture. Mice were euthanized at experimental endpoint by isoflurane anesthesia followed by carbon dioxide inhalation. Bacterial burden was determined by collecting colon samples into 1 mL of reduced PBS and homogenized in a FastPrep-24 (MP Biomedicals) at 5.5 m/s for 2 minutes. Sample homogenate was diluted for plating on MacConkey agar (Difco) and incubated for 18-20 hours at 37 °C before counting bacterial colonies.

Chemically-induced colitis was triggered by exposure to 3% dextran sulfate sodium (DSS) in the drinking water for 4 days before being placed on regular drinking water. Mice were sacrificed at 5 days post-treatment onset.

Colonization with both a-virulent *C. rodentium* Δ*escN* and commensal *E. coli* Mt1B1 were carried out by oral gavage with 2.5-4×10^8^ CFU from overnight culture, as determined by retrospective plating. Control WT-infected mice were inoculated at the same dosage. *S.* Typhimurium (ATCC 14028) infections were carried out by oral gavage with 1×10^6^ CFU of *S.* Typhimurium from overnight culture. Fecal samples were processed for CFU enumeration as described above.

### Bacterial growth at physiological pH

Bacterial growth at physiological pH was determined by inoculating at an Optical Density at 600 nm (OD600) of 0.005 from an overnight culture into 250 µL total volume of the media of interest (LB or DMEM base media adjusted to physiological pH). Cultures were incubated at 37 °C with agitation for 20 hours in a Synergy H1 plate reader (Biotek) with OD600 measurements at 10-minute intervals.

### CMT-93 cell culture assays

Murine rectal carcinoma cell line CMT-93 (ATCC CCL-223) was maintained in DMEM supplemented with 10% fetal bovine serum, 1% Glutamax, and 1% non-essential amino acids. Cells were used between passages 4-10. Cells were seeded in 12-well plates at 90,000 cells/well and incubated for 48 hours prior to infection. Confluent CMT-93 cells were washed once with phosphate buffered saline (PBS) before infection. The *C. rodentium* inoculum was prepared by sub-culturing 1:40 from overnight culture to mid-log phase (3.5 hours, 37 °C, shaking) in either LB or DMEM media adjusted to intestinal pH. Subcultures were washed and resuspended in cell culture media at multiplicity of infection (MOI) 100. Infected plates were centrifuged (1000 rpm, 5 minutes) to synchronize the infection before incubation for 4 hours at 37 °C and 5% CO2. Infected plates were washed 5 times with PBS before detachment using 0.1% Triton X-100 in PBS for dilution and plating on neutral LB agar. Adherent bacterial CFU were quantified and data was normalized to exact inoculum dosage (determined by retrospective plating) to account for differences between between pH conditions.

### Bacterial and eukaryotic gene expression analysis

*C. rodentium* was subcultured for 3.5 hours 1:40 in LB base media adjusted to gastrointestinal pH, as before. After subculture samples were preserved in RNAprotect Bacterial Reagent (Qiagen) and stored at ™70 °C until extraction. Bacterial RNA extractions were done using the RNAse-Free DNase Set (Cat. no. 79254, Qiagen) as per the manufacturer’s instructions. A QuantiTect Reverse Transcription Kit (Cat. no. 205313, Qiagen) was used for cDNA synthesis and Quantitative RT-qPCR was performed using a QuantiNova SYBR Green RT-PCR Kit (Cat. no. 208056) in combination with primers (Table S2) for target gene mRNA. *dnaQ* was used as an endogenous control. Data was normalized to efficiency of primers by target gene (91.6-103.5% for all primers used).

Eukaryotic RNA was isolated from mouse tissue samples (stomach and colon) stored in RNAprotect Tissue Reagent (Qiagen) at ™70 °C before extraction using a GeneJET RNA Purification Kit (Cat. no. K0731, Thermo Scientific). cDNA synthesis and Quantitative RT-qPCR was performed as described above using various primer pairs (Table S2). β-2- microglobulin (*B2M*) was used as an endogenous control.

### Bacterial cell morphology

*C.* rodentium was cultured to mid-log phase (3.5 hours, 37 °C with agitation) at physiological pH before being spun down and resuspended in PBS with the addition of FM 1-43 Dye (*N*-(3-Triethylammoniumpropyl)-4-(4-(Dibutylamino) Styryl) Pyridinium Dibromide (Cat. no. T3163, Invitrogen). Images were taken on a Zeiss X10 light microscope and cell area was measured using CellProfiler 4.2.1 (33,34).

### Bacterial survival assay for ATR response

*C. rodentium* was cultured overnight at either neutral pH 7 or colonic pH 5.7 under anaerobic conditions (90% N2, 5% CO2, 5% H2 at 37 °C without agitation). Overnight cultures were diluted 1:20 into unadjusted LB (the baseline “untreated” condition), or LB adjusted to gastric pH 3.5. At 30-minute intervals, subcultures were sampled and diluted in PBS for plating on neutral LB agar to determine viability by quantifying surviving CFU.

### Intra-bacterial redox potential

Assays were performed using a protocol adapted from (35,36). In brief, a roGFP2 *C. rodentium* reporter strain was grown overnight in LB containing 100 μg/mL of carbenicillin. Overnight cultures were diluted 1:25 into 50 mL of pre-warmed LB without antibiotics and were grown for 5 hours at 37 °C in a shaking incubator. Bacterial strains were washed and resuspended in saline solution (0.9% NaCl w/w). Cultures were then exposed to the different pH of interest in a black 96-well plate with a volume of 200 µL and an OD600 of 1. Intra-bacterial redox was assessed by measuring fluorescence with an excitation at 405 and 480 nm and an emission at 510 nm. Background fluorescence was controlled by measuring the fluorescence of WT *C. rodentium*. Normalized 405/480-nm ratios were calculated by using a fully oxidized (100 mM H2O2) and reduced (10 mM DTT) control conditions.

### ELISA analysis of fasting serum gastrin levels

Blood was collected from naïve and day 8- infected mice following a 12-hour fast. Whole blood was allowed to clot for 2 hours on ice before serum collection. Gastrin levels were analyzed using the mouse gastrin enzyme-linked immunosorbent assay (ELISA) kit (Novus Biologicals, Cat. No. NBP3-08148). Absorbance at 450 nm was measured using a Synergy H1 plate reader (Biotek).

### Statistical analysis

Statistical analysis was performed in Graphpad Prism (www.graphpad.com) and clarified in figure legends. Unless otherwise stated, analysis was performed using a Mann-Whitney test to compare two groups and a one-way ANOVA with Šidák’s multiple comparisons test for more than two groups. Aggregate results represent the mean +/- SD, and statistical significance is represented by \**p* <0.05, \*\**p* <0.01, \*\*\**p* <0.001 and \*\*\*\**p* <0.0001.

## Results

### Regional variability in gastrointestinal pH may be influenced by genetic background

To assess the response of *C. rodentium* to intestinal pH, we first defined regional pH ranges within its natural host, the mouse. We compared reported gastrointestinal pH (pH 2-6) to pH measured in healthy, female, 7-week-old C57BL/6 mice across gut regions (Figure 1A-B). We observed a similar pattern to reported murine and human values, which see pH increase at significant increments towards the ileum-large intestinal regions (Figure 1B). At the cecum, we found a non-significant drop in pH which is characteristic of human intestinal pH, but not previously observed in mice (3). Across all regions, we observed variability between individuals, particularly in the stomach and ileum, demonstrating that physiological pH likely fluctuates over time and with varied consumption of food and water.

Overall however, the pH across both the small intestine (duodenum, jejunum, ileum) and large intestine (cecum, colon) was more basic than reported murine values (6). Mice in the aforementioned study were fed a different diet, the Teklad Global 18% Protein Rodent Diet (Teklad), with a lower protein content (18%) (37). Dietary protein is thought to increase intestinal pH by increasing ammonia production as a by-product of bacterial metabolism (4,11,38). We therefore placed C57BL/6 mice on either the Teklad diet or Picolab Rodent Diet 20 5053 (Picolab; 21% protein) for two weeks before sacrifice. We found no significant differences between diets, except in the distal colon where pH was significantly higher on the lower protein Teklad diet (Figure S1A-B). This indicates that the more alkaline pH observed across gut regions may instead be due to differences in mouse genetic background (BALB/c vs C57BL/6).

To investigate whether mouse genetic background can indeed result in altered intestinal pH, we compared the intestinal pH of both BALB/c and C57BL/6 mice fed the Picolab diet. We found a significant difference in intestinal pH in the ileum and colon regions (Figure S1C). As reported previously, we did not observe a drop in intestinal pH of BALB/c mice from the ileum to cecum regions, rather pH increased along the length of the GI tract. This indicates that genetic differences between mouse strains may result in altered intestinal pH, with pH of C57BL/6 mice more closely reflecting the pattern found in humans. Altogether, these data establish important physiological variation within the naive murine gastrointestinal tract, which could be used by enteric pathogens to signal their biogeographical location.

### *C. rodentium* alters growth dynamics, epithelial attachment, and virulence gene expression in response to gastrointestinal pH

Having established the range of physiologically relevant pH, we next investigated whether *C. rodentium* responds to variations in pH by adjusting the pH of LB to mimic the ranges of pH found in both mouse and human GI tracts. Growth of *C. rodentium* was highest at the range of pH 6.4-7.5, with impaired growth at acidic pHs 4.4 and 4.8, resembling the gastric and duodenal regions (Figure 2A). As previously reported (27), no growth was observed at gastric pH of 3.5-4. Human enteropathogens EHEC, EPEC, and *S.* Typhimurium were similarly unable to grow at gastric pH 3.5, and displayed altered growth dynamics in response to pH below 5 (Figure S2A-C). This suggests that while acid tolerance and viability rates differ between pathogens (30,39), they are all affected by fluctuations in gastrointestinal pH.

**Figure 2.**
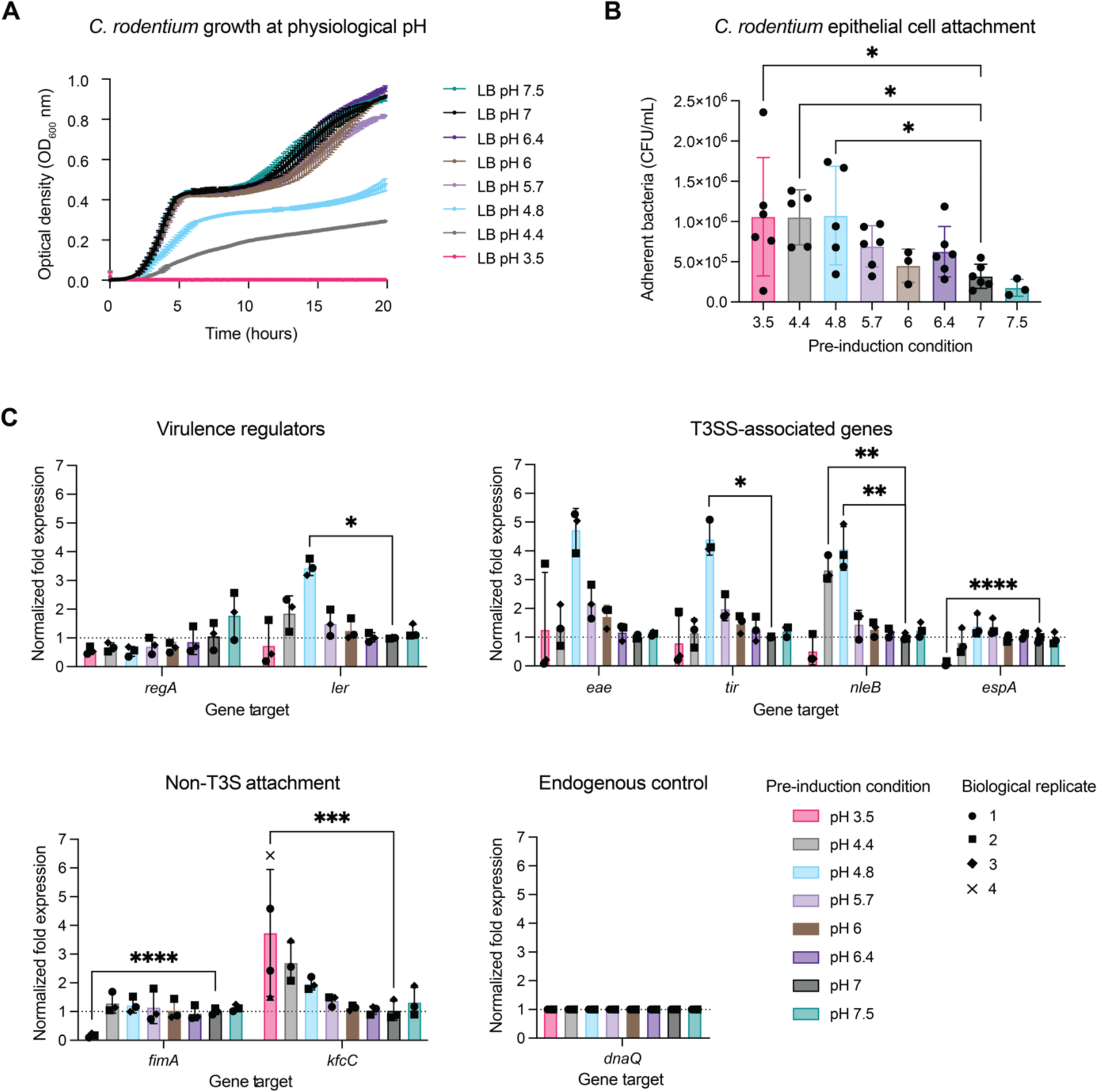
*C. rodentium* alters its behaviour and virulence in response to regional physiological pH. A) Growth in pH-adjusted LB across the range of gastrointestinal pH, measured as optical density (OD) 600 nm at 10-minute intervals over 20 hours. B) Attachment of *C. rodentium* to CMT-93 murine colonic epithelial cells after pre-induction at gastrointestinal pH. Data represent biological replicates (N = 3-4; each the average of three technical replicates). Statistical analysis represents an ordinary one-way ANOVA test with Holm-Šidák’s multiple comparisons test. C) Gene expression analysis by RT-qPCR of key virulence- and early infection-related genes across the range of physiological pH. Statistical analysis represents t-tests with Bonferroni correction for multiple comparisons. Error represents mean +/- SD.

We further investigated the relative metabolic activity of *C. rodentium* subjected to intestinal pH by measuring the reduction of MTT (3-(4,5-dimethylthiazol-2-yl)-2,5- diphenyltetrazolium bromide) to formazan by NADPH-dependent oxidoreductase enzymes in metabolically active bacteria. As expected, bacteria incubated at pH 3.5 produced significantly less formazan compared to neutral pH despite an increased biomass, suggesting decreased metabolic activity (Figure S2D-E). These results support both the absence of growth at pH 3.5 and the low recoverability of cells after removal from acidic pH (27). The observed increase in biomass at pH 3.5 may further suggest the active encapsulation of community members within biofilms, an important stress-induced protective behaviour (40). Taken together, extreme low pH of the gastric environment has considerable consequences to bacterial fitness.

To determine if the altered fitness dynamics observed across physiological pH also reflect differences in pathogen virulence, we assessed epithelial cell attachment, an essential characteristic of infection. This was done by ‘pre-inducing’ *C. rodentium* via subculture in pH-adjusted LB before infection of CMT-93 cells, a murine rectal cell line chosen because the rectum is a known large intestinal niche of *C. rodentium* during host colonization. We expected attachment to increase in response to pH representing key replicative niches of *C. rodentium*, such as the cecum and colon. However, *C. rodentium* attachment was significantly increased after pre-induction at pH 3.5 and 4.4, representing gastric-duodenal pH (Figure 2B). Attachment was lowest at pH 7.5 which represents the upper-range of ileal pH, as well as pH found in the extra-intestinal environment. Assays were repeated using subculture in pH-adjusted DMEM media, which is known to activate expression of the T3SS. Despite high buffering of DMEM media making accuracy difficult, the same trends were noted in both growth and attachment (Figure S2F-G). Therefore, LB media was used in all subsequent assays.

To further explore pH-mediated virulence regulation we characterized the gene expression profile of known virulence- and early infection-associated genes following *C. rodentium* subculture in pH-adjusted LB prior to RNA extraction. Despite the fact that LB media is not known to activate the T3SS, type III secretion-associated virulence genes *ler, tir, eae,* and effector gene *nleB*, were highly expressed at pH 4.8 which represents duodenal pH (a 4-5 fold increase from baseline). At gastric pH 3.5 we observed down-regulation of virulence genes associated with type III secretion, including both structural and effector-associated genes (Figure 2C), as well as virulence regulators *ler* and *regA* (41). While a known fimbrial gene *fimA* was also down-regulated, we found an upregulation of *kfcC*, a component of the *C. rodentium* type IV pilus, which may explain the increase in epithelial cell attachment observed at gastric pH (41).

To further investigate the extent to which the T3SS contributes to pH-mediated attachment, we evaluated the ability of a Δ*escN C. rodentium* strain lacking the T3SS to attach to CMT-93 cells following pre-induction as before. We found that infection with the Δ*escN* strain did not result in decreased epithelial attachment following pre-induction in LB at low pH 3.5 and 4.4 (Figure S2H), further supporting that low pH induced epithelial attachment is mediated by non-T3SS factors. Together, these data demonstrate that *C. rodentium* regulates virulence gene expression in a pH-dependent manner, leading to increased attachment-related behaviour upon initial entry to the intestinal environment.

### Intestinal pH activates transcription of diverse bacterial stress responses

Bacterial stress responses and virulence are closely co-regulated (30,42). Therefore, we next investigated the relationship between intestinal pH and the transcriptional expression of stress-related genes. A key strategy of food-borne pathogens under acid stress is to increase intracellular pH, which can be done via production of amino acid decarboxylases which allow for the consumption of cytoplasmic protons (43). Interestingly, we did not find homology within the *C. rodentium* genome to either the transcriptional regulators GadW and GadX of the glutamate decarboxylase acid-resistance system, nor to AdiC of the arginine acid-resistance system found in some *E. coli* strains, including EHEC (44,45). We did however identify 70% protein sequence homology between the *E. coli* CadA, a lysine decarboxylase, and *C. rodentium* LdcC (Figure S3). As expected, we found that transcript levels of *ldcC* increased at low pH from 4.4-5.7, indicating that *C. rodentium* also uses amino acid decarboxylation to combat acid-induced stress despite lacking homology to two major systems (Figure 3A).

**Figure 3.**
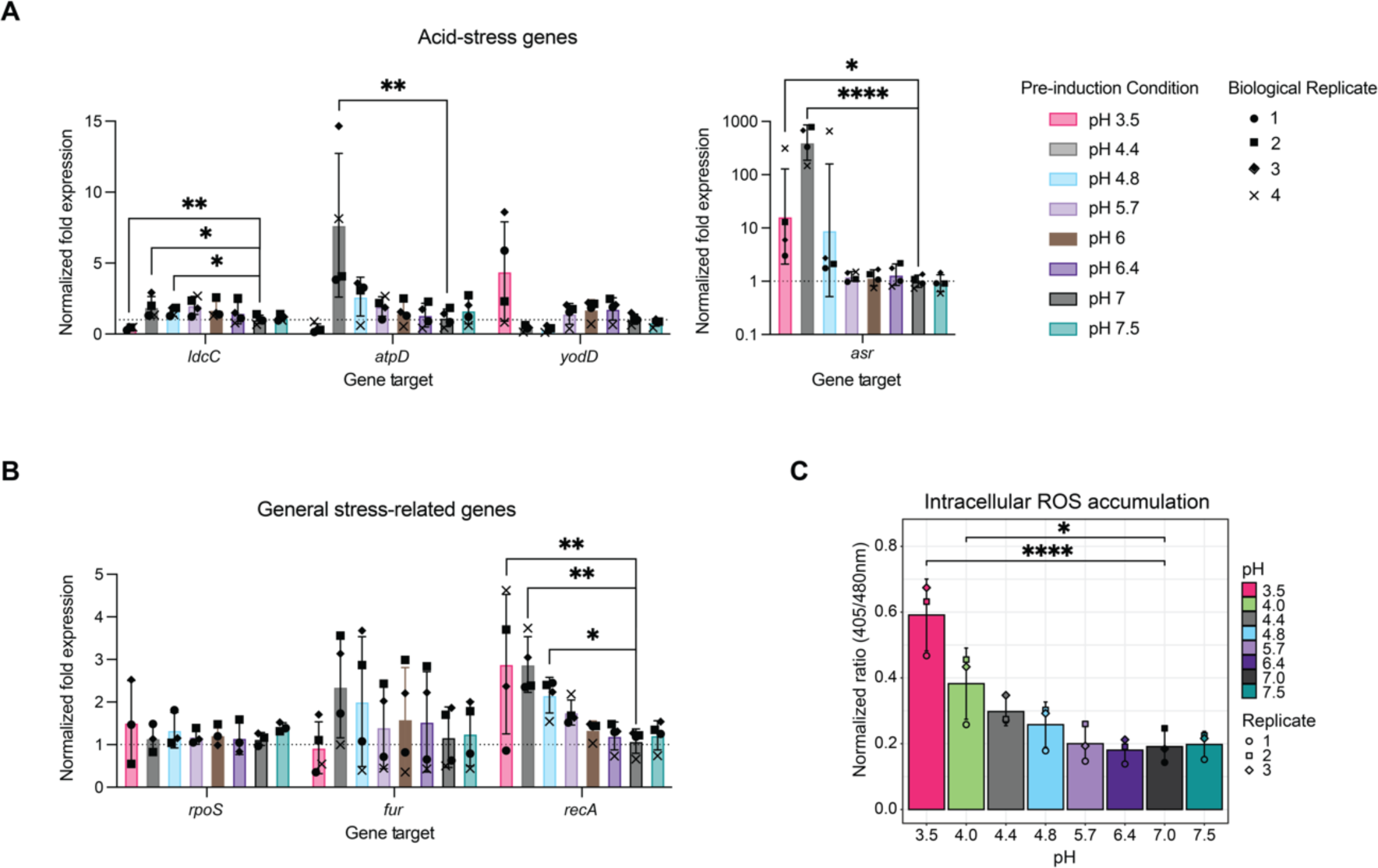
Stress gene regulation is affected by physiological pH. A) Expression of acid stress-associated genes across the range of gastrointestinal pH by qPCR. Data represent biological replicates (average of three technical replicates; N=4). B) Expression of general stress-associated genes across the range of gastrointestinal pH by qPCR. Data represent biological replicates (average of three technical replicates; N=3-4). C) Intracellular reactive oxygen species (ROS) accumulation at 3 hours post-inoculation into LB at physiological pH, as determined using a rhoGFP reporter *C. rodentium* strain. Data represent biological replicates (average of three technical replicates; N=3). Statistical analysis represents t-tests with Bonferroni correction for multiple comparisons.

Another strategy of food-borne pathogens in dealing with acid-stress is proton efflux (43). The F1F0-ATPase proton efflux membrane ATPase is conserved across many bacterial pathogens, including *C. rodentium*, in which it is encoded by the *atpD* gene. We observed a 7- fold increase in expression of *atpD* at pH 4.4 compared to neutral media (Figure 3A). Additionally, upon searching the *C. rodentium* genome for genes annotated as involved in acid resistance, we chose to further investigate the expression of *yodD*, a stress-induced peroxide/acid resistance protein, and *asr*, the acid resistance repetitive basic protein, both of which are shared with EHEC. Both genes were upregulated at gastric pH 3.5, and *asr* in particular demonstrated a 300-fold increase in expression at pH 4.4, suggesting that both of these genes are important to *Citrobacter*’s acid resistance strategy within the upper GI tract.

Aside from direct mechanisms of acid resistance, we further profiled the effect of intestinal pH on non-specific stress responses which have evolved as general mechanisms to overcome environmental stress. We investigated the expression of *rpoS*, a gene encoding the stationary phase sigma factor present in Gram-negative bacteria which controls general stress responses to stressors such as oxidative or osmotic stress. RpoS is important for *C. rodentium* survival under heat and H2O2 conditions, as well as full pathogen virulence (46,47). Mutation of *rpoS* in EHEC has also been found to be decrease resistance to acid stress (48). While *rpoS* expression was therefore expected to increase with decreased pH, it was unaltered across the pH range tested (Figure 3B). This was surprising, given that when we exposed a roGFP2 reporter strain of *C. rodentium*, which can be used to measure intra-bacterial redox dynamics, to different pHs we found that gastric pH 3.5-4 significantly increased oxidative stress (Figure 3C) (36). We further investigated expression of *fur*, encoding a ferric iron transcription regulator which may regulate transcription of genes that protect against ROS damage (43). While we observed a trend towards increased *fur* expression at pH 4.4, there was no significant up- or down-regulation of the system induced by altered pH. Despite the lack of ROS-specific gene expression, we did observe evidence that bacteria at low pH respond to general cell damage. For example, we found that expression of *recA* which encodes a recombinase protein used to repair DNA double-strand breaks increased with increased acidity of the culture media, suggesting cell stress and DNA damage at pH 4.8 and lower (Figure 3B) (49).

### pH-induced stress alters membrane regulation and cellular morphology

An important outcome of stress-induced gene regulation is the repair and maintenance of the bacterial cell membrane, and can even induce adaptive changes to cellular morphology. To visualize potential morphological changes induced by pH, *C. rodentium* was pre-induced at GI pH before staining with FM 1-43 Dye. We observed bacterial cell lengthening at pH 4.0-4.8, an indicator of cell stress, and a trend towards small size and abnormal cell shape at pH 3.5 (Figure 4A-B). We also observed a decrease in membrane intensity in a sub-population of cells at pH 3.5, possibly reflecting a loss of membrane integrity (27).

**Figure 4.**
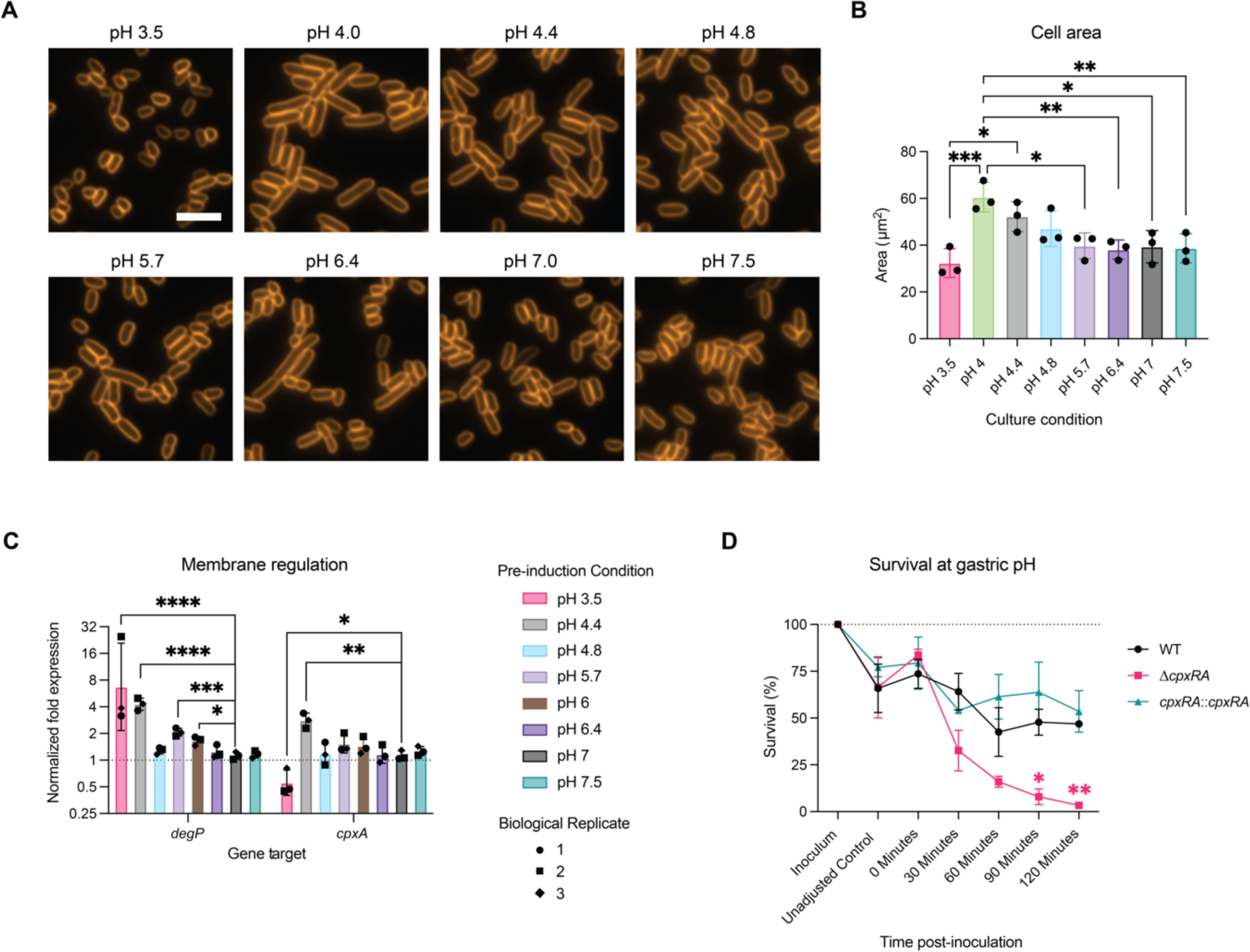
Membrane regulation in response to intestinal pH. A) Representative cellular morphology after 3.5 hours at relevant physiological pHs by light microscopy and cell membrane staining. Scale bar represents 5 µm. B) Cell area measurements from light microscopy across physiological pH (N=3 biological replicates, each the sum of 2 technical replicates). C) Expression of genes by qPCR associated with the cpx stress response following culture at gastrointestinal pH. Statistical analysis represents t-tests with Bonferroni correction for multiple comparisons. Data represent biological replicates (average of three technical replicates; N=3). D) Survival of wild-type, Δ*cpxRA,* cpxRA::cpxRA *C. rodentium* in gastric pH 3.5 over the range of average gastric emptying time. Each point represents the average of 3-5 biological replicates (3 technical replicates per biological replicate). Statistical analysis represents a mixed-effects model with Geisser-Greenhouse correction and Šidák’s multiple comparisons test. Colour of stars indicates significance of Δ*cpx* (pink) or complement (green) strains compared to the WT strain.

The cpx envelope stress response is an important mechanism to overcome conditions which result in misfolding of envelope proteins, such as alkaline pH, and has been shown to be required for *C. rodentium* virulence (42,50). We found that *cpxA*, part of the CpxRA two-component system which mediates the cpx response, was most highly expressed at low pH 4.4 (42). CpxRA is known to regulate both the expression of periplasmic proteases such as DegP, as well as T3SS-related virulence. We found that expression of *degP* which is induced by stress (51), was significantly upregulated in response to subculture at pH 3.5, 4.4, 5.7, and 6.0 compared to neutral base media (Figure 4C). *degP* was not upregulated at pH 4.8 which resulted in a strong activation of T3SS-related genes, consistent with reported inverse regulation of these systems (Figure 2C), and supporting cpx stress response regulation under low upper intestinal pH conditions.

Successful gut transit must require fluid regulation of membrane protein synthesis and degradation in response to intestinal pH. We therefore tested survival at gastric pH 3.5 of wild-type *C. rodentium* or a strain deficient in the CpxRA two-component system (Δ*cpxRA*), and which is unable to properly regulate its membrane stress response (42). While there was no difference upon initial exposure to pH 3.5 (0 minutes) between wild-type, Δ*cpxRA*, and Δ*cpxRA::cpxRA* strains, the Δ*cpxRA* strain exhibited significantly reduced survival after 30-120 minutes, with up to an average of 3.4% viability in the absence of *cpxRA* after 120 minutes, which represents the upper limit of gastric emptying time, as compared to 53.5% in the complement strain (Figure 4D) (52). The cpx response is therefore not only necessary for *C. rodentium* virulence, but is also critical to gastric passage and overall host-host transmission. Overall, these data suggest that *C. rodentium* alters its membrane and overall shape under physiological pH conditions to adapt to its surrounding environment.

### Host-adaptation increases *C. rodentium* acid tolerance which may support host transmission

*C. rodentium* is known to undergo host-adaptation during gastrointestinal passage, allowsing for more effective transmission to new hosts at a reduced infectious dose (53,54). We hypothesized that a crucial part of this host-adapted (HA) state would include adaptation to better tolerate the gastric environment. While the conditions that trigger host-adaptation remain unknown, since host-adaptation occurs following passage through the mouse gut, we investigated whether *in vitro* culture at colonic pH representing the key *in vivo* niche of *C. rodentium*, ahead of gastric pH exposure, could induce an acid tolerance response (ATR) and mimic the protection observed after gastrointestinal transit. Indeed, we observed a striking ATR following stationary phase growth at colonic pH 5.7 under anaerobic conditions (Figure 5A). While culture at neutral pH resulted in only 20.6% survival after two hours at gastric pH, subculture at colonic pH 5.7 resulted in an average survival rate of 83.6%. Anaerobic conditions were chosen both to mimic passage through the low oxygen large intestinal environment (55). We did not observe an ATR under aerobic conditions, nor in response to short-term exposure to low pH (pre-induction to mid-log phase) (Figure S4A).

**Figure 5.**
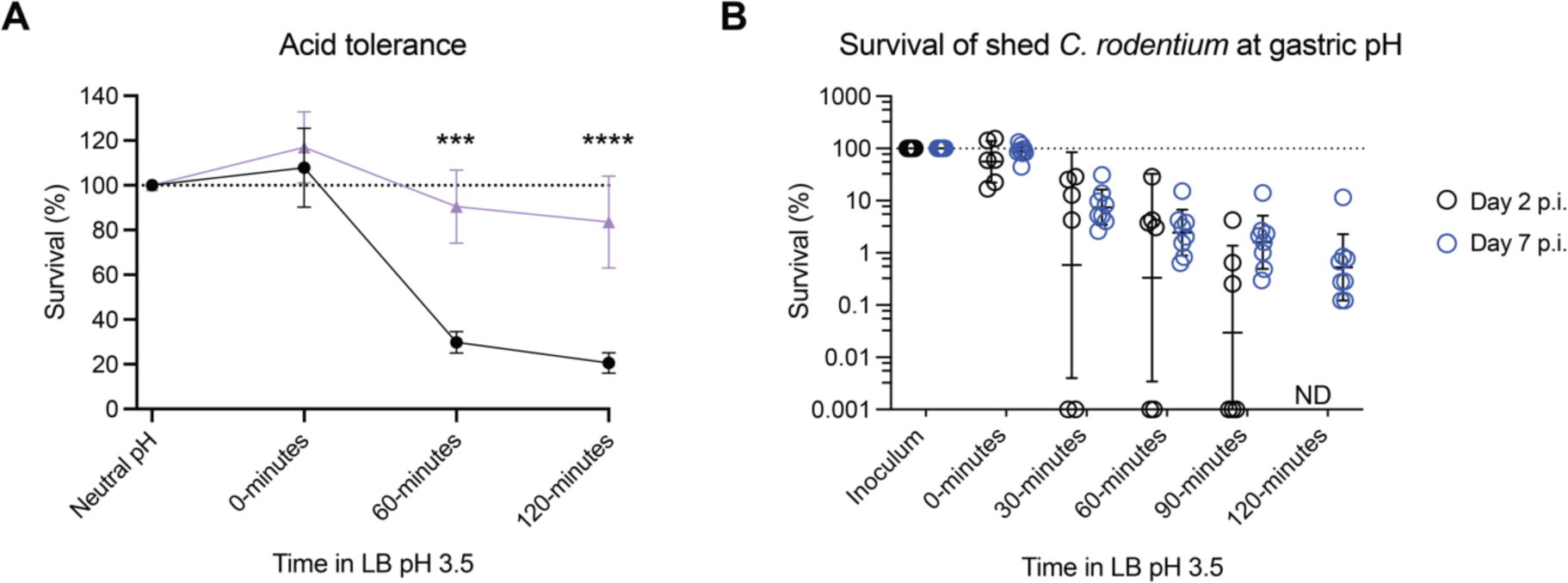
Host-adapted *C. rodentium* is better able to survive acid-induced stress. A) *C. rodentium* survival at gastric pH after initial anaerobic pre-induction at either colonic pH 5.7 or neutral pH. Each point represents the average of 3 biological replicates (3 technical replicates per biological replicate). Statistics represent a two-way ANOVA with Šidák’s multiple comparisons test. B) Survival of *C. rodentium* in homogenized feces collected on days 2 and 7 post-infection, representing non-adapted and host-adapted states (N=6-8). ND, not detected.

Having demonstrated a robust ATR under colonic pH conditions, we next sought to determine whether HA-*C. rodentium* conditioned within the host gut, demonstrates increased acid tolerance. We evaluated the survival of *C. rodentium* shed in the feces of infected mice at day 2, representing an early, non-HA state, and *C. rodentium* shed at day 7 post-infection, representing the host-adapted state (Figure 5B). Though we found similar percent survival at 0- 30 minutes post-exposure to pH 3.5, none of the day 2-shed *C. rodentium* was able to survive 120 minutes at gastric pH. As these fecal samples were homogenized before exposure to pH 3.5 rather than naturally encapsulated in the fecal pellet, we repeated this assay on day 2 feces without homogenization. Despite an increase to 8% survival, we still observed significant population loss (Figure S4B). This indicates that while natural encapsulation, such as within contaminated food, contributes to ‘naïve’ non-HA pathogen passage of the gastric barrier it must be combined with acid adaptation to facilitate high efficiency transmission. Altogether, these data suggest that *C. rodentium* adapts to the acidic conditions of the colon during colonization, possibly conferring a competitive advantage to initial gut seeding.

### Changes to host cell regulation and circulating gastrin levels result in decreased gastric pH following *C. rodentium* infection

All the above experiments were done by subjecting *C. rodentium* to the range of intestinal pH found within the naïve murine gut. However, considering the global gut environmental changes that occur during enteric infection, we expected changes to intestinal pH post-infection (pi), either as a result of pathogen colonization or an active host response. Therefore, we measured gastric and cecal pH of infected mice at days 4 and 8 pi and compared them to uninfected controls. Cecal pH did not change at days 4 or 8 pi, despite a high pathogen burden (Figure 6A). However, we found that gastric pH significantly decreased by day 8 pi as compared to uninfected mice, correlating with colonic pathogen burden (Figure 6A-B). This was not due to decreased food consumption by the mice (Figure S5A-B) (6).

**Figure 6.**
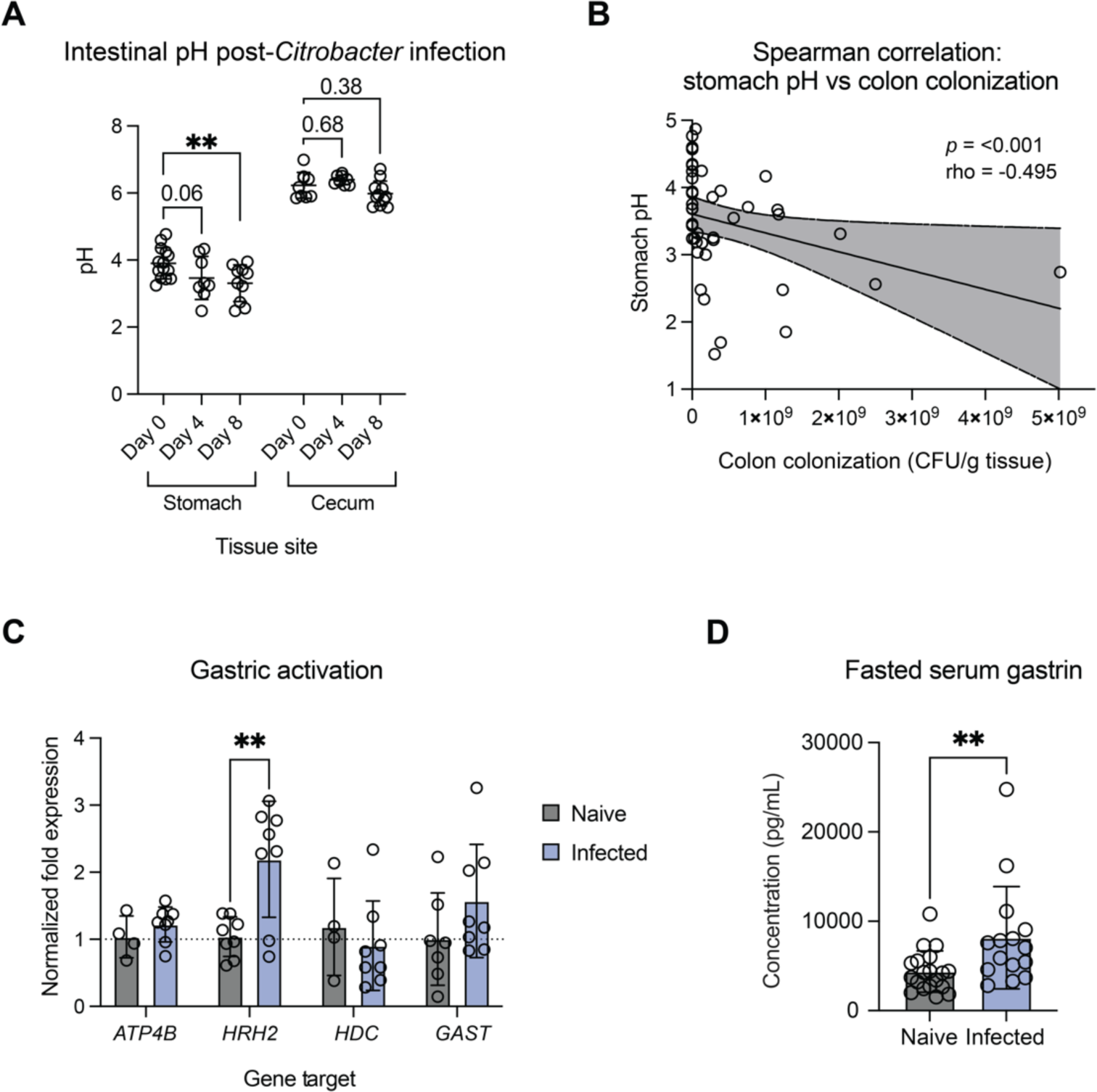
Murine gastric pH decreases at peak infection. A) Intestinal pH measurements in the murine stomach and cecum over time post-*C. rodentium* infection (inoculum dose of 10^8^ CFU). Statistical analysis represents an Ordinary two-way ANOVA with Dunnett’s multiple comparisons test (N = 8-14). B) Spearman correlation of stomach content pH and colon colonization (CFU/gram tissue) at Days 4 and 8 post-infection. Line represents linear regression +/- 95% confidence interval. *P*- and rho-values represent Spearman correlation values. C) Gene expression analysis of host genes associated with stomach parietal cell activation and acid secretion within stomach tissue from naïve and infected mice (N = 4-8). Statistics represent a two-way ANOVA with Šidák’s multiple comparisons test. D) Fasted serum gastrin levels in naïve and day 8- infected mice. Statistical analysis represents an unpaired t-test (N = 15-19).

To determine whether this decrease in gastric pH was the result of altered acid secretion, we analyzed host gene expression within the stomach tissue of naïve and infected mice at day 8 pi. We observed a non-significant increase in *ATP4B* which encodes the beta subunit of the gastric H+,K+-ATPase, a proton pump responsible for acid secretion in parietal cells (Figure 6C) (56). We then specifically investigated changes across histamine-, gastrin-, and acetylcholine-mediated pathways of gastric cell activation. We observed a significant increase in *HRH2* expression, encoding histamine receptor H2, a G protein-coupled receptor that stimulates gastric acid secretion in response to histamine (Figure 6C) (56–58). Additional genes associated with histamine-mediated acid secretion, such as *HDC* (encoding the histidine decarboxylase which converts L-histidine to histamine), and *PAC1* (encoding a receptor involved in histamine release from gastric enterochromaffin-like cells) remained unaltered (Figures 6C and S5C).

Hormonal activation of gastric acid secretion is facilitated by release of gastrin, a hormone produced by gastric G cells and released into the circulation following food consumption (56–58). We observed a non-significant trend towards increased *GAST* gene expression, which directly encodes gastrin (Figure 6C). Expression of *CCK2R*, encoding the cholecystokinin B receptor which responds to gastrin, was highly variable in expression among infected mice (Figure S5C). Furthermore, although cholinergic agents also facilitate parietal cell activation, we did not observe changes to the expression of *CHRM3*, a gene encoding the M3 muscarinic acetylcholine receptor which responds to the release of cholinergic agents (Figure S5C).

We also investigated genes related to negative control of gastric acid secretion, specifically gastric levels of *VIP* gene expression encoding vasoactive intestinal peptide and the gene encoding its associated receptor in the stomach, *VPAC2* (56–58). We found a non-significant trend towards decreased levels of *VPAC2* (Figure S5D; *p* = 0.08), indicating that infection potentially leads to dysregulation of multiple systems involved in maintaining gastric pH. Collectively, these data suggest changes to parietal cell activation post-infection.

To assess whether transcript-level expression reflects systemic changes to regulation of gastric pH, we measured fasting serum gastrin levels by ELISA. High levels of circulating gastrin detected in a fasted state are indicative of hypergastrinemia, and gastrin levels are thereby used as a diagnostic marker in humans (57). Indeed, we found that circulating gastrin levels in *C. rodentium*-infected mice at day 8 pi were significantly higher than in naïve mice (Figure 6D). Taken together, our data support decreased gastric pH post-*C. rodentium* infection, resulting from active changes to host acid regulation.

### Gastric pH decreases in response to further infection and chemical models of epithelial barrier dysfunction

To further investigate decreased gastric pH as an active host defense mechanism, we next wished to determine whether similar changes could be observed in other enteric infection models. To address this, we infected mice with human pathogen *Salmonella enterica* serovar Typhimurium and measured gastric pH at peak infection on day 4 pi. We observed a significant decrease in gastric pH in response to *S.* Typhimurium (Figure 7A) which significantly correlated with pathogen burden (Figure S6A), demonstrating consistency across additional enteric infection models. To determine whether this response is infection-specific, we next performed a dextran sodium sulfate (DSS) model of chemically induced colitis. Despite no difference between the pH of control and DSS drinking water (Figure S6B), we again observed a significant decrease in gastric pH following 4 days of chemical administration (Figure 7A). We therefore hypothesized that gastric pH may decrease in response to intestinal damage rather than infection specifically.

**Figure 7.**
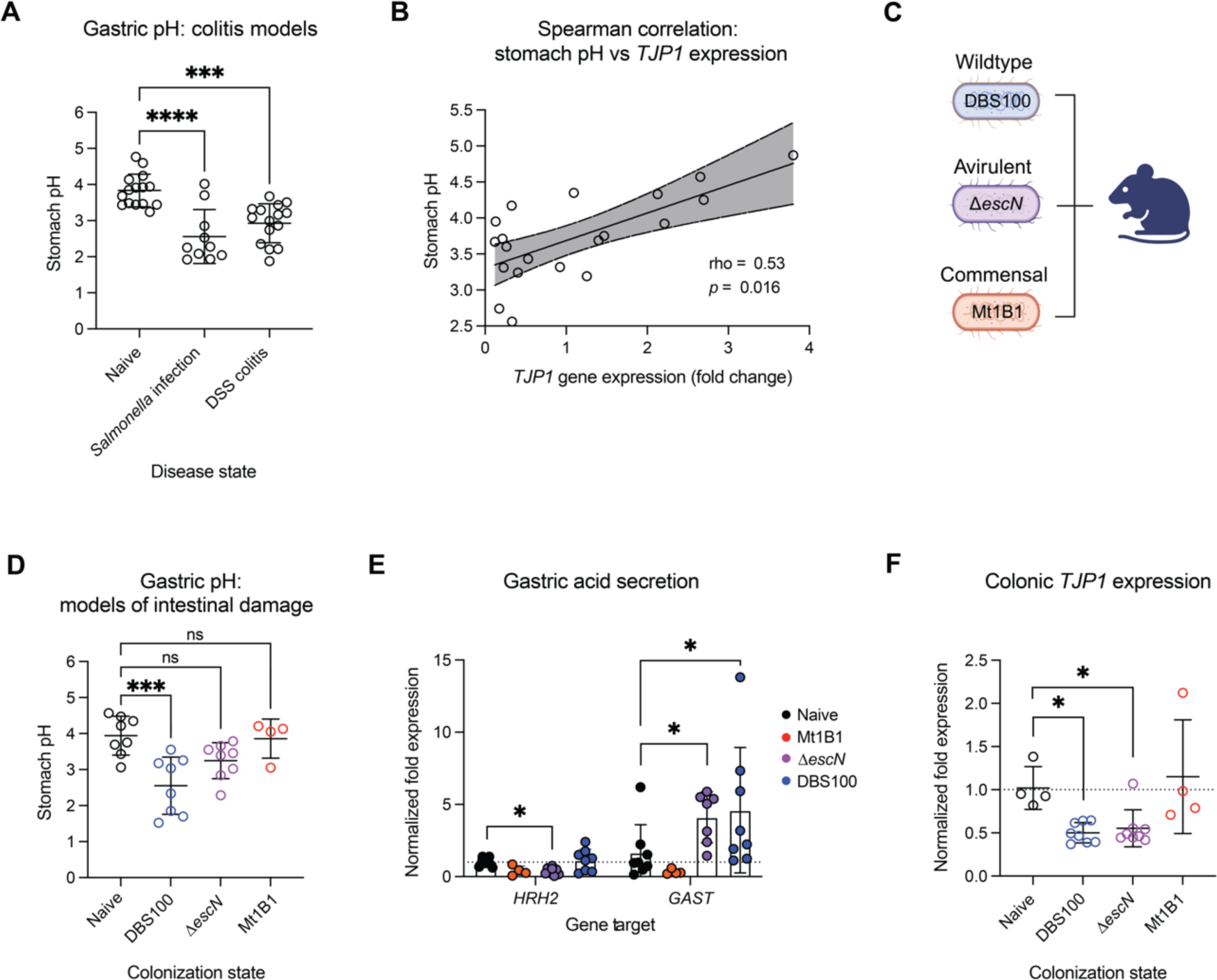
Murine gastric pH decreases in additional models of intestinal damage. A) Measurements of gastric pH in additional models of intestinal damage: *Salmonella enterica* serovar Typhimurium typhoid model of infection (*Salmonella* infection), and the dextran sulfate sodium model of colitis (DSS colitis) (N = 10-15). Statistical analysis represents an ordinary one-way ANOVA with Tukey’s multiple comparisons test. B) Spearman correlation of stomach pH and *TJP1* expression within colonic tissues of naïve and infected mice (N = 20). Line represents linear regression +/- 95% confidence interval. *P*- and rho-values represent Spearman correlation values. C) Summary of strains used to test the impact of intestinal damage on gastric pH: wild-type *C. rodentium* DBS100, T3SS-deficient Δ*escN C. rodentium*, and commensal *E. coli* Mt1B1. D) Measurement of gastric pH in mice which are uninfected, or colonized with either wild-type *C. rodentium*, Δ*escN C. rodentium*, or commensal isolate *E. coli* mt1B1 at an inoculum dose of 2-4×10^9^ CFU. Measurements were taken at sacrifice at day 4 post-inoculation (N = 4-8). Statistical analysis represents an Ordinary one-way ANOVA with Bonferroni’s multiple comparisons test. E) Gene expression analysis of host genes *HRH2* and *GAST* within stomach tissue from naïve mice and mice colonized with either wild-type *C. rodentium*, Δ*escN C. rodentium*, or commensal isolate *E. coli* mt1B1 at day 4 post-inoculation (N = 4-8). Statistics represent a two-way ANOVA with Šidák’s multiple comparisons test. F) Colonic *TJP1* expression in naïve mice and mice colonized with either wild-type *C. rodentium*, Δ*escN C. rodentium*, or commensal isolate *E. coli* mt1B1 at day 4 post-inoculation (N = 4-8). Statistics represent a Kruskal-Wallis test with Dunn’s multiple comparisons test.

To investigate the relationship between intestinal damage and gastric pH, we measured colonic expression of tight junction protein *TJP1*, a marker of epithelial barrier integrity. Indeed, we observed a positive correlation between *TJP1* expression and gastric pH during *C. rodentium* infection (rho = 0.53, *p* = 0.02; Figure 7B; Figure S6C), such that lower gastric pH reflected decreased colonic barrier integrity. In contrast, serum levels of calprotectin, a general marker of systemic host inflammatory state altered by infection, did not correlate with gastric pH (rho = - 0.299, *p* = 0.2; Figure S6D-E).

To further test the relationship between intestinal damage and gastric pH, we colonized mice with either wild-type *C. rodentium*, a type III secretion system-deficient mutant strain (*ΔescN*), or murine commensal *E. coli* isolate Mt1B1 (Figure 7C). The Δ*escN* and commensal Mt1B1 strains are capable of causing only a moderate to negligible level of tissue pathology and inflammation upon murine colonization, respectively (59,60), as confirmed by inflammatory cytokine levels in the colon (Figure S6F). Due to the inability of the Δ*escN* to persist within the mouse gut, we sampled mice at 4 days pi when bacterial burdens were expected to be similar across all groups. We found that gastric pH did not decrease in response to commensal *E. coli* Mt1B1, and that Δ*escN* inoculation resulted in only a mild, non-significant, decrease in gastric pH (Figure 7D). Furthermore, unlike in wild-type *C. rodentium* infection, colonization rates of Δ*escN* DBS100 and Mt1B1 did not correlate with gastric pH (Figure S6G), suggesting that bacterial colonization in the absence of tissue pathology does not affect gastric pH.

We further measured expression levels of *HRH2*, which were found to be significantly lower in response to colonization by the Δ*escN* strain, with no differences observed between naïve and Mt1B1-colonized mice (Figure 7E). However, we did find elevated expression of *GAST* in Δ*escN* colonized mice, likely accounting for the moderate decrease observed in gastric pH (Figure 7E). This increase in *GAST* expression again corresponded with decreased expression of *TJP1* in the colon of mice colonized with the Δ*escN* strain but not *E. coli* Mt1B1, further supporting the relationship between colonic epithelial barrier integrity and gastric pH (Figure 7F). Together, these results indicate that decreased gastric pH may be a widespread host response to intestinal damage.

## Discussion

Gastrointestinal pH varies along both the human and mouse gut, likely serving as an important indicator of intestinal geography for invading pathogens. Our study reveals that enteric pathogen *C. rodentium* responds to subtle pH changes representative of gut pH (even changes of < 0.5), altering its growth and virulence. Exposure to pH resembling the cecal environment, *C. rodentium*’s optimal niche, promoted pathogen growth but did not stimulate the expected T3SS- related attachment. Instead, we found that small intestinal pH 4.4-5.7 increased expression of T3SS genes. Duodenal pH may signal pathogen entry to the small intestine from the acidic stomach environment, a transition that necessitates attachment to the host epithelium to prevent pathogen expulsion from the GI tract. Interestingly, gastric pH also enhanced epithelial cell attachment, this time through upregulation of the type IV pilus Kfc (K99 fimbrial-like adhesin), an early attachment factor, rather than the T3SS (Figure 2). Collectively, our data suggest that low pH is a trigger of loose intestinal attachment upon immediate entry to the small intestine, while the higher pH of the large bowel favors pathogen expansion to support the a high pathogen load (27).

Unexpectedly, we found that gastric pH was significantly decreased at peak infection. Meanwhile, downstream intestinal pH did not change, despite high pathogen burdens (Figure 6). Upregulation of host genes associated with acid secretion suggests that stomach acidification is an active adaptation of the host. Histamine receptor-2 (*HRH2*) in particular is associated with gastric ulcers such that HRH2-blockers, such as rantidine (Zantac), are used to treat conditions involving excessive stomach acid production (61,62). We found that this response may not be pathogen-specific, but instead related to intestinal damage, demonstrating the same phenotype during chemically induced colitis but not in response to colonization by commensal *E. coli* nor by Δ*escN C. rodentium* lacking key damage-inducing virulence machinery (Figure 7). The host environment has evolved many adaptative responses to clear invading bacteria such as fever, vomiting, and both specific and non-specific immune responses (1,63). Decreasing stomach pH may be another important host response for protection against further pathogen ingestion.

While a decrease from an average gastric pH of 4.0 to 3.5 during peak *C. rodentium* infection may seem inconsequential, we observed significant differences in cellular morphology and metabolic activity between pHs. While pH 3.5 induced small cell size and decreased metabolic activity, a pH increase of only 0.5 resulted in cell elongation and metabolic recovery, and a reduction in intracellular ROS (Figures 3-4; Figure S2). Therefore, even incremental host changes may present a formidable barrier to intestinal re-entry. Interestingly, ‘host-adapted’ *C. rodentium* shed at peak infection demonstrated higher acid tolerance than host-naïve *C. rodentium*, indicating that the pathogen develops a more robust acid tolerance response during passage through the mouse gut, which may be triggered by exposure to colonic pH conditions (Figure 5). Acid adaptation may thereby contribute to the enhanced transmission of host-adapted *C. rodentium* as compared to host-naïve *C. rodentium* (53). Taken together, it is clear that adaptation occurs on both sides to maintain the balance between pathogen and host.

Given the impact of enteric infection on gastric acid secretion in mice, intestinal damage and inflammatory state may have important functional consequences to gut pH homeostasis. Few studies investigate human gastrointestinal pH following infection or other stressors. However, a known association exists between functional dyspepsia (FD) and infectious gastrointestinal disease by bacteria, viruses, and protozoa (64). FD is a disorder of unknown etiology characterized by stomach pain and other indigestive symptoms, which occasionally overlaps with post-infectious IBS (65,66). A meta-analysis of 17 clinical studies found onset of post-infectious FD in 9.5% of adult subjects, indicating a 2.5-fold risk of FD following acute gastroenteritis (64). Therefore, further understanding of gastrointestinal pH in response to environmental triggers or altered host health, such as infectious disease state, could be important to predicting susceptibility to secondary intestinal disease.

In summary, our study illustrates how regional GI fluctuations in pH signal changes to *C. rodentium* behaviour and virulence, with consequences to bacterial stress and survival. We further observed infection-induced alterations to host pH which could impact the bioavailability and effectiveness of pH-dependent release drugs, or colonization of probiotic bacterial strains for the treatment of intestinal morbidities. As such, there is immense value in understanding the complex and long-lasting interactions of pathogens and their hosts.

## Acknowledgements

The authors thank all of our colleagues in the Finlay laboratory for their support and assistance, especially W Deng, L Thorson, and T Bozorgmehr. This work was supported by grants from the Canadian Institutes of Health Research (CIHR) to BB Finlay (FDN-159935). SE Woodward is a CIHR CGS-D Graduate Scholar (GSD-154171) and was supported by a UBC Four Year Fellowship and Dmitry Apel Memorial Scholarship. Supporting images were created with BioRender.com.

## Author Contributions

S.E.W. conceived the project. S.E.W. and L.M.N. designed experiments. S.E.W., L.M.N., J.P-D., W.F., A.S.P., and I.T. performed experiments and analyzed data. S.E.W. and A.S.P. performed animal work. S.E.W. wrote the original draft of the manuscript with input from all authors. All authors revised the manuscript. B.B.F. acquired funding for the project and provided supervision.

