## Supplemental Materials for "Pathogen and host adapt pH responses during enteric infection"

Supplemental Methods

**Analysis of rodent diets.** To compare standard rodent diets, mice were fed either the Teklad Global 18% protein diet (Cat. no. 2918; Envigo) or Picolab Rodent Diet 20 (Cat. no. 5053; LabDiet), over a period of two weeks before measurement. Mouse chow pH was measured by dissolving pellets in 3 mL deionized water per 100 mg.

**Biofilm formation and metabolic activity.** To assess pH-induced biofilm formation, *C. rodentium* was inoculated in LB at physiological pH to an OD_600_ of 0.5 in a black, clear bottom 96-well plate. The plate was incubated at 37 °C, standing, for 48 hours to allow for biofilm formation. After 48 hours, culture media was removed and the OD_600_ of the resultant biofilm was measured in a Synergy H1 plate reader (Biotek). 100 µL of 3-(4,5-dimethylthiazol-2-yl)-2,5-diphenyltetrazolium bromide (MTT) at a concentration of 0.5 mg/mL in LB was added to the biofilms and allowed to incubate at 37 °C for 10 minutes. MTT was removed and replaced with 100 µL of 100 % DMSO for resuspension before measuring OD at 570 nm to approximate formazan production as a readout of metabolic activity.

**ELISA analysis of inflammation.** Serum calprotectin levels were analyzed using the mouse S100A9 ELISA kit (Hycult), as per the manufacturer’s instructions. Serum was collected from non-fasted mice by allowing whole blood to clot for 2 hours on ice before spinning to collect serum. Absorbance at 450 nm was measured using a Synergy H1 plate reader (Biotek).

**Cytokine measurements.** To analyze intestinal inflammation of mice colonized with either *E.* coli Mt1B1, ∆*escN* *C. rodentium*, or WT *C. rodentium* as compared to naïve animals, colon tissue samples were collected at 4 days post-inoculation by first removing intestinal contents. Samples were collected into 1mL of PBS containing a protease inhibitor cocktail (Roche Diagnostics) and homogenized using a FastPrep-24 (MP Biomedicals) at 5.5 m/s for 2 min. Homogenate was then spun down at 16,000 x *g* for 20 minutes at 4 ºC for supernatant collection. Inflammatory cytokines were measured using a mouse inflammation cytometric bead array (CBA) kit (BD Biosciences, Cat. No. 552364) according to manufacturer’s instructions. Measurements were made on an Attune NxT flow cytometer (Thermo Fisher Scientifice), normalizing cytokine concentrations to tissue weight.

Supplemental Figures and Tables

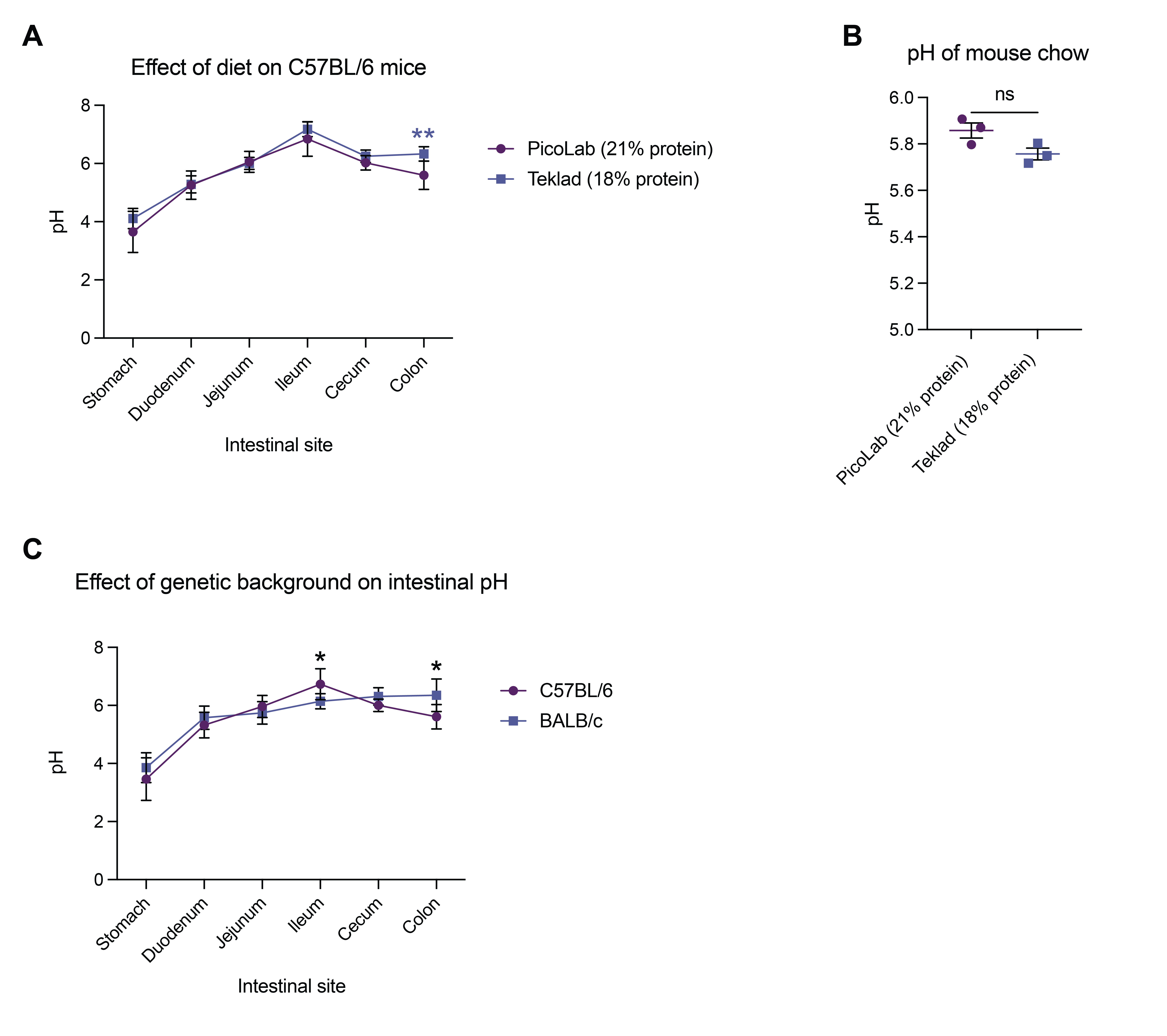

Figure S1. Effect of standard mouse chow on intestinal pH of C57BL/6 mice. A) Murine intestinal pH after two weeks on either standard Teklad 18% protein mouse chow or Picolab 21% protein mouse chow. Statistical analysis represents a mixed-effects model with Geisser-Greenhouse correction for matched values and Tukey’s multiple comparisons test (N = 6). B) pH of Teklad 18% protein standard mouse chow compared to Picolab 21% protein standard mouse chow. Statistical analysis represents an unpaired t-test (N = 3 pellets, average of 3 pH readings). C) Murine intestinal pH of C57BL/6 and BALB/c mice after two weeks on Picolab 21 % protein mouse chow. Statistical analysis represents a mixed-effects model with Geisser-Greenhouse correction for matched values and Tukey’s multiple comparisons test (N = 8-18).

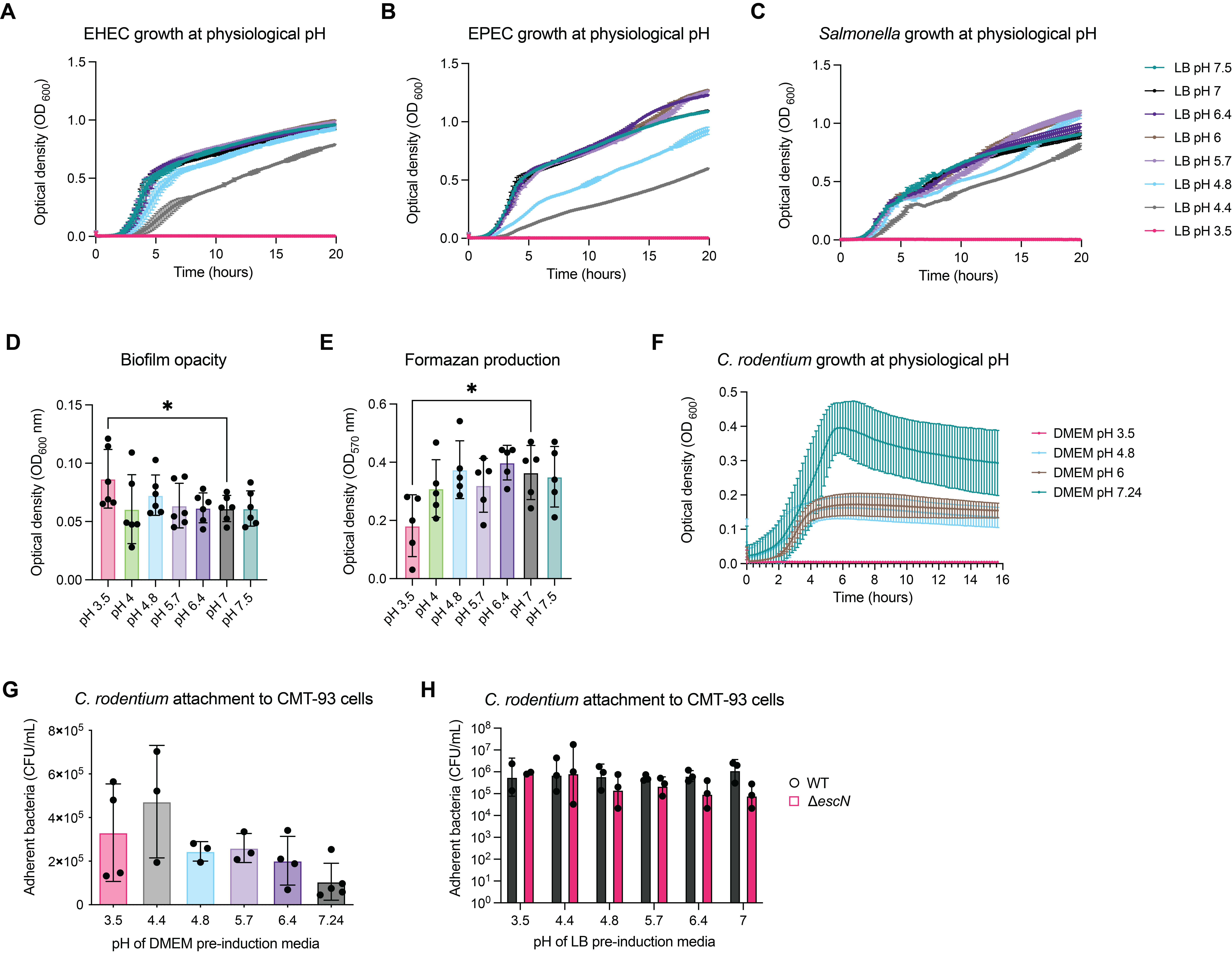

Figure S2. Figure S2. Similar patterns of growth observed in human pathogens EHEC, EPEC, and *S.* Typhimurium at physiological pH. Growth of A) enterohemorrhagic *E. coli* (EHEC), B) enteropathogenic *E. coli* (EPEC), and C) *S.* Typhimurium at physiological pH over 20 hours, indicating that all three pathogens are unable to grow at pH 3.5. Data represent 3 biological replicates (the average of 3 technical replicates). Error represents mean +/- SD. D) Biofilm opacity measured by optical density at 600 nm after 48 hours of static growth seeded at physiological pH (N = 6, each the sum of 3 technical replicates). Statistical analysis represents a Friedman test with Dunn’s multiple comparisons test. E) Metabolic activity within the biofilm after 48 hours of static growth seeded at physiological pH, measured by the production of formazan from MTT at OD 570 nm. Error represents mean +/- SD. F) Growth of *C. rodentium* at physiological pH in DMEM base media over 20 hours. Data represent 3 biological replicates (the average of 3 technical replicates). Error represents mean +/- SD. G) Attachment of *C. rodentium* to CMT-93 murine colonic epithelial cells after pre-induction in DMEM base media adjusted to gastrointestinal pH. Data represent biological replicates (the average of 3 technical replicates). H) Attachment of wild-type (WT) or T3SS-deficient (∆*escN*) *C. rodentium* to colonic epithelial cells after pre-induction at gastrointestinal pH. Data represent biological replicates (the average of 3 technical replicates).

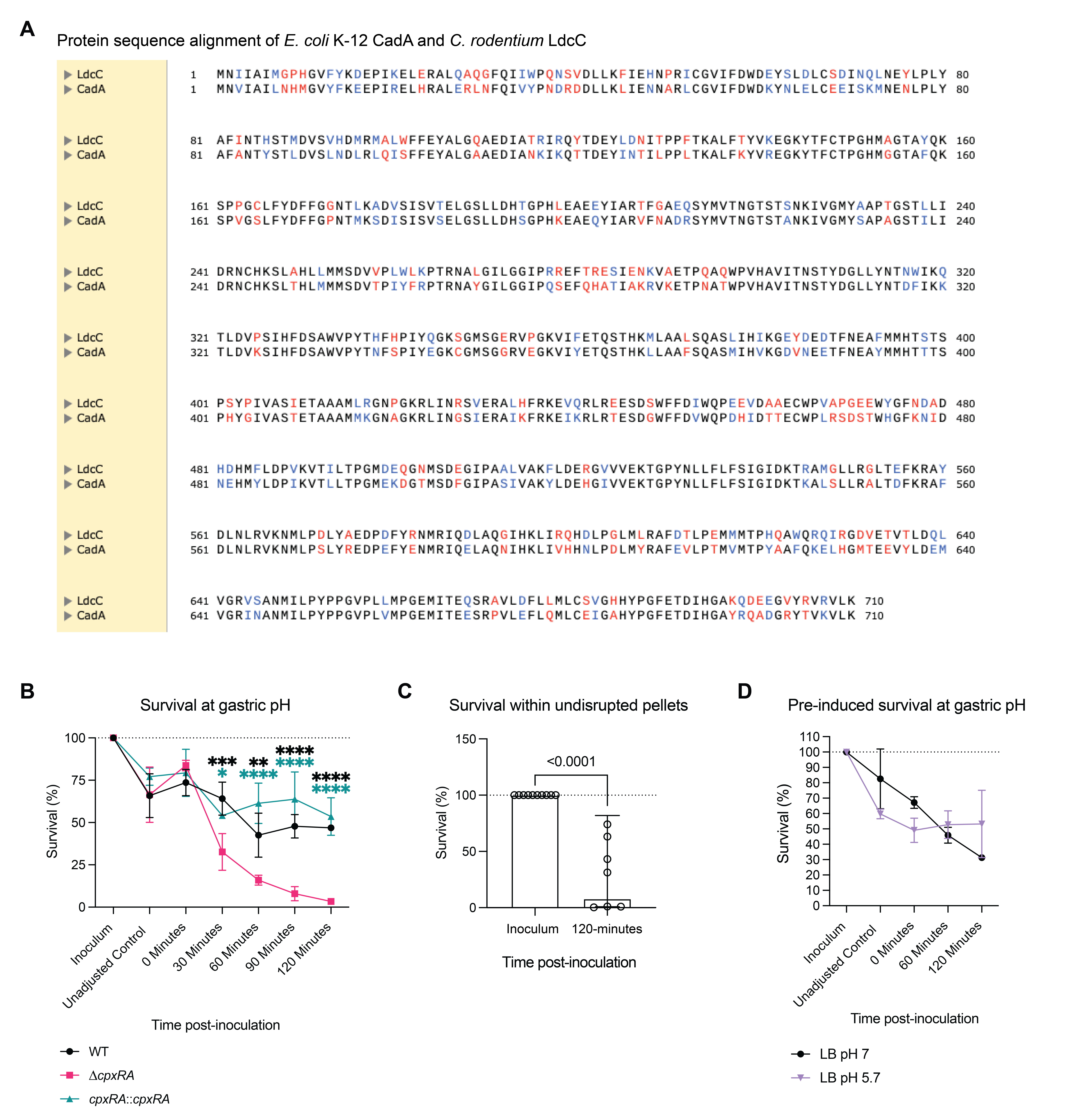

Figure S3. Homology to the *E. coli* CadA lysine decarboxylase. A) Protein sequence homology between *C. rodentium* LdcC and *E. coli* CadA.

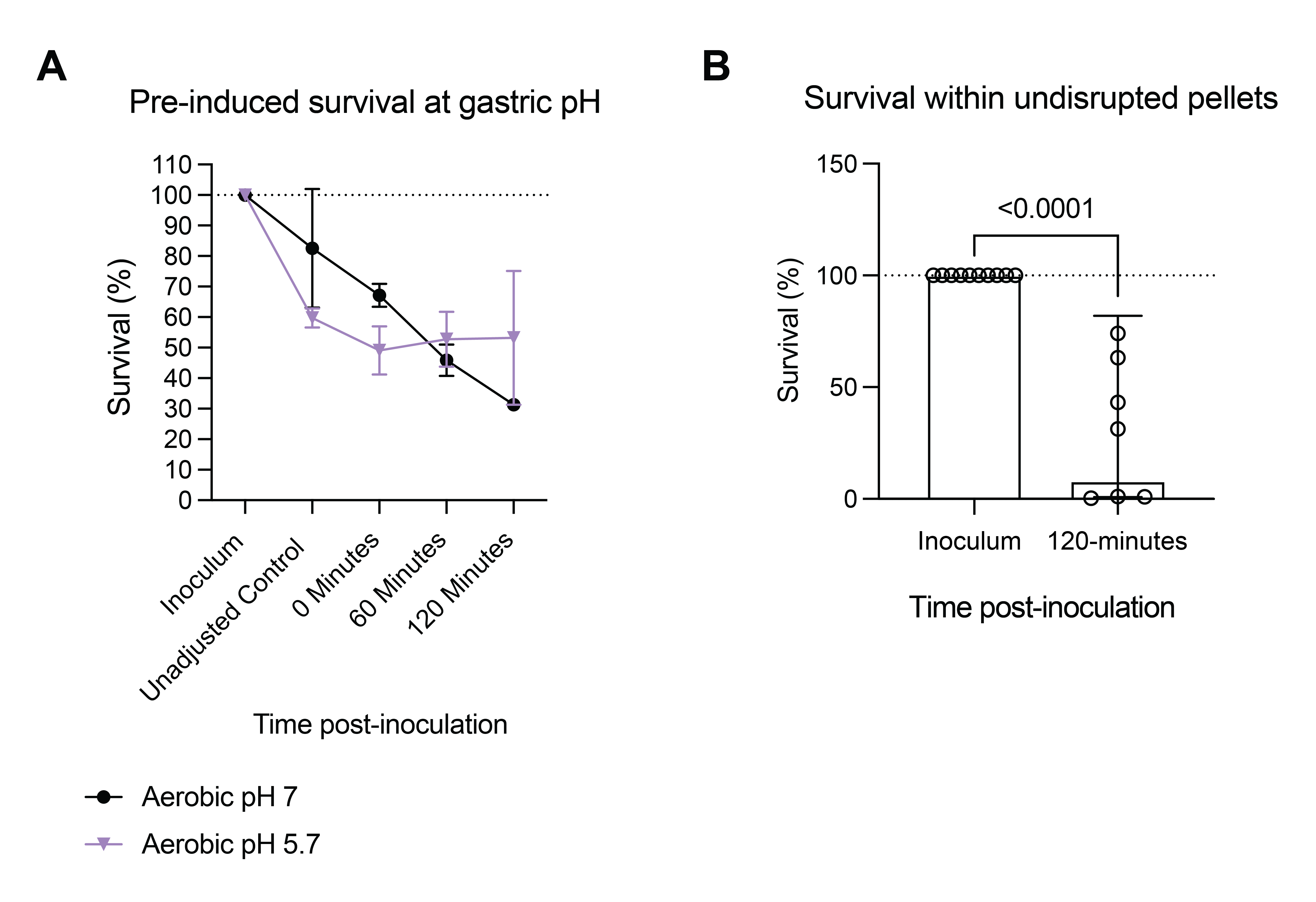

Figure S4. *C. rodentium* survival at gastric pH. A) Survival of *C. rodentium* pre-induced aerobically at either neutral pH 7 or colonic pH 5.7 for 3.5 hours (to mid-log phase) before exposure to gastric pH 3.5. Each point represents the average of 3 biological replicates (3 technical replicates per biological replicate). Statistics represent a mixed-effects model with Geisser-Greenhouse correction and Šidák’s multiple comparisons test. Error represents mean +/- SD. B) Survival of *C. rodentium* shed in the feces on Day 2 post-infection (pi) at gastric pH 3.5. Fecal pellets were not disrupted before submersion in LB pH 3.5. Statistical analysis represents a Mann-Whitney test (N = 6-7).

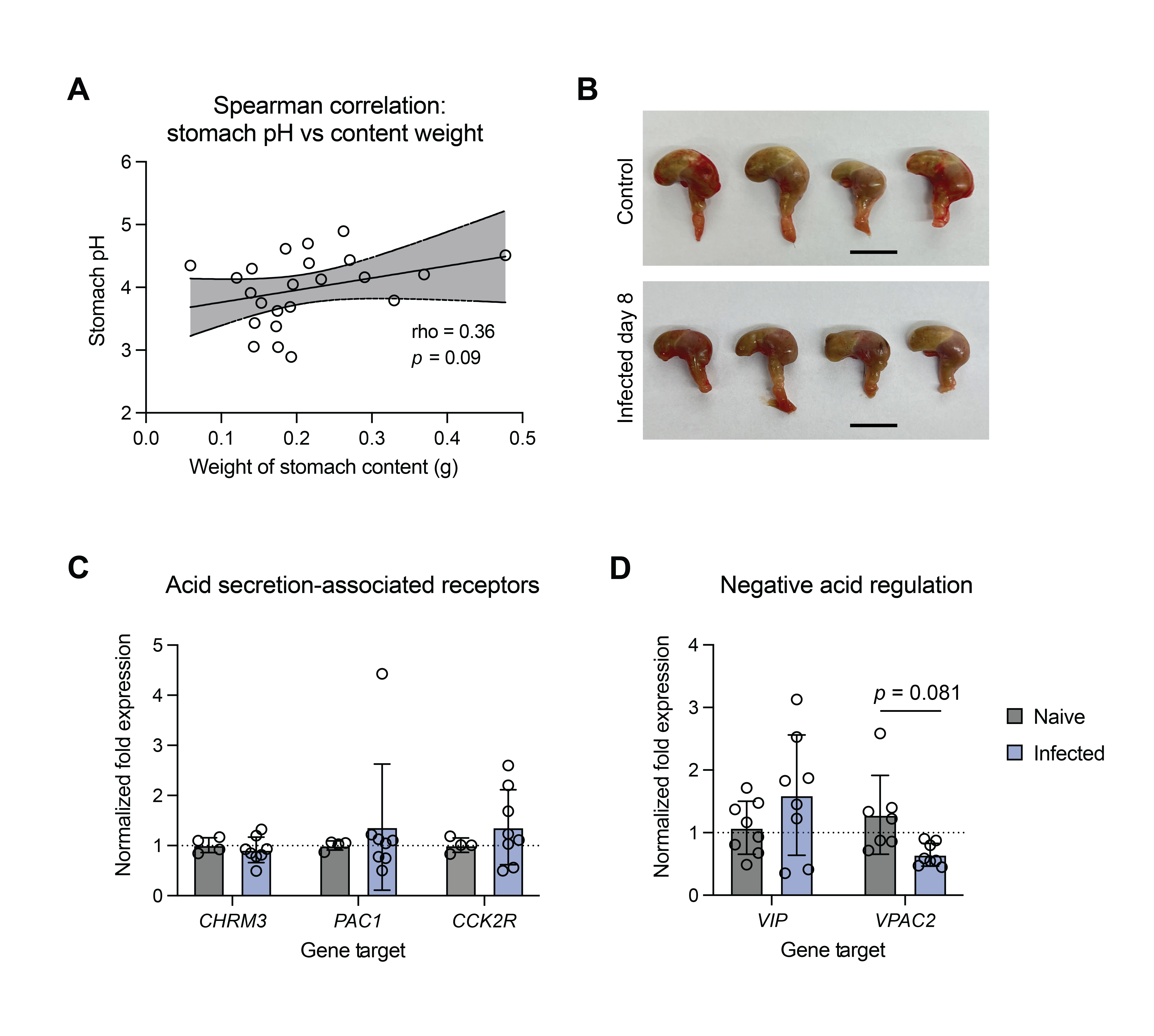

Figure S5. Factors involved in gastric acid secretion post-infection. A) Spearman correlation of stomach pH and weight of stomach contents (N = 23). Line represents linear regression +/- 95% confidence interval. *p*- and rho-values represent Spearman correlation values. B) Representative stomach images from fasted control and day-8 infected mice. Scale bar represents 1 cm. C) Normalized fold expression of genes associated with host regulation of acid secretion within stomach tissue from naïve and infected mice (N = 4-8). Statistics represent a two-way ANOVA with Šidák’s multiple comparisons test. D) Normalized fold expression of genes associated with host negative regulation of acid secretion within stomach tissue from naïve and infected mice (N = 7-8). Statistics represent a two-way ANOVA with Šidák’s multiple comparisons test.

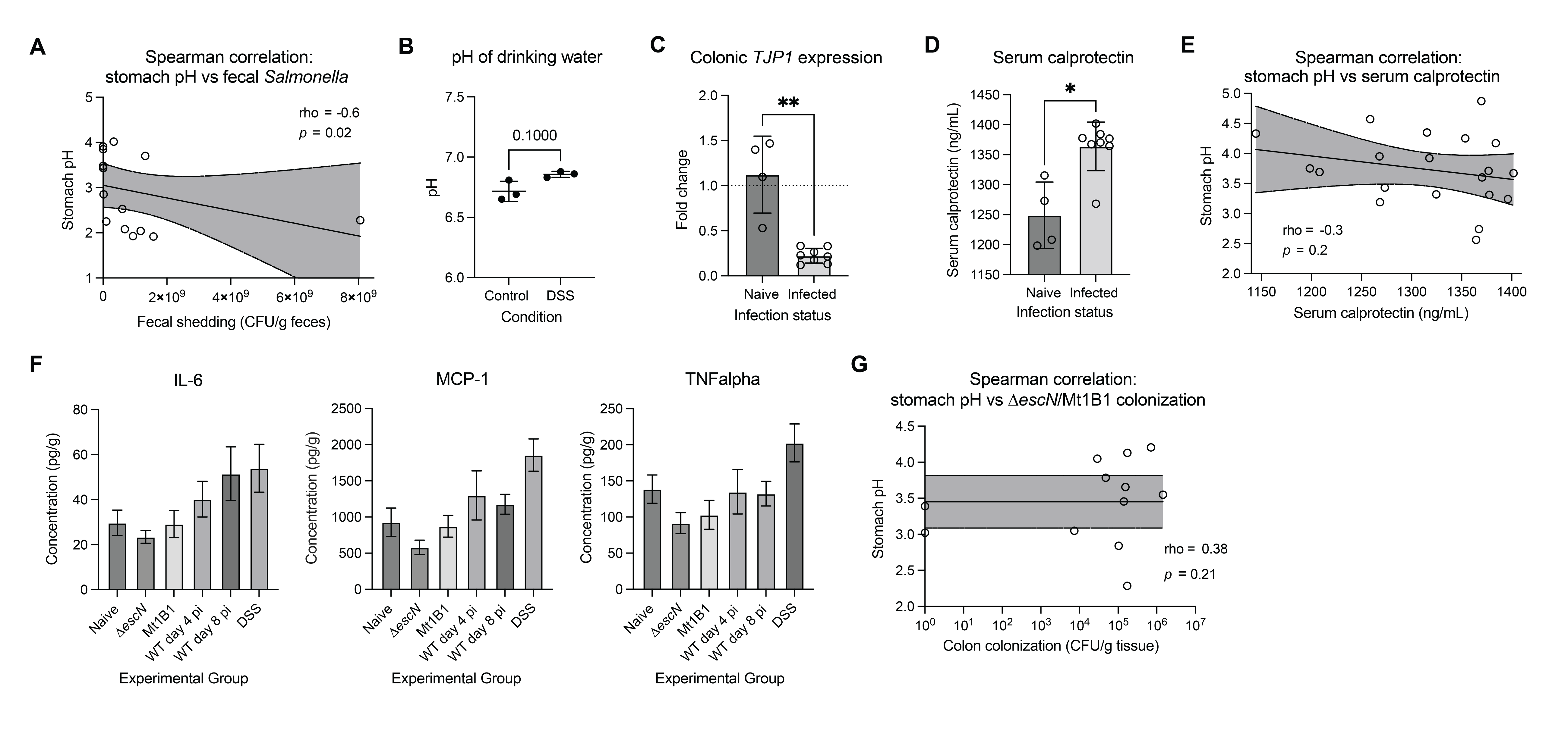

Figure S6. The relationship of stomach pH with infection, epithelial barrier integrity, and inflammation. A) Spearman correlation of stomach pH and fecal burden of *Salmonella* Typhimurium at day 4 of a murine typhoid infection model (N = 14). Line represents linear regression +/- 95% confidence interval. *p*- and rho-values represent Spearman correlation values. B) pH of drinking water used in the DSS-colitis model. C) Colonic expression of *TJP1* in naïve and *C. rodentium*-infected mice at day 8 post-infection (N = 4-8). Statistics represent a Mann-Whitney test. D) Serum calprotectin levels measured by ELISA in naïve and *C. rodentium*-infected mice at day 8 post-infection (N = 4-8). Statistics represent a Mann-Whitney test. E) Spearman correlation of stomach pH and serum calprotectin levels of naïve and infected mice (N = 20). Line represents linear regression +/- 95% confidence interval. *p*- and rho-values represent Spearman correlation values. F) Cytokine levels measured in the colons of naïve mice or mice colonized with ∆*escN* *C. rodentium*, commensal *E. coli* Mt1B1, or wild-type (WT) *C. rodentium* at day 4 post-inoculation, or WT *C. rodentium* at day 8 post-infection, or mice treated with dextran sodium sulfate (DSS) to induce colitis (N = 4-8). G) Spearman correlation of colonization levels by avirulent ∆*escN* *C. rodentium* and commensal isolate *E. coli* Mt1B1 with stomach pH (N = 10). Line represents linear regression +/- 95% confidence interval. *p*- and rho-values represent Spearman correlation values.

Table S1. Bacterial strains used in this study.

| **Strain** | **Description** | **Source or reference** |
| --- | --- | --- |
| DBS100 | Wild-type *C. rodentium*; ATCC 51459 | (Schauer et al., 1995) |
| EDL933 | Wild-type EHEC originally isolated from ground beef; ATCC 43895 | (Riley et al., 1983) |
| E2348/69 | Wild-type EPEC | (Levine et al., 1978) |
| CDC 6516-60 | Wild-type *S.* Typhimurium; ATCC 14028 |  |
| DBS100 ∆*cpxRA* | *C. rodentium* lacking genes encoding the cpxRA two-component system | (Thomassin et al., 2015) |
| DBS100 ∆*escN* | *C. rodentium* unable to produce the type III secretion system | (Gauthier et al., 2003) |
| Mt1B1 | Commensal *E. coli* isolated from murine gut | (Serapio-Palacios et al., 2022) |

Table S2. Oligonucleotide primers used for RT-qPCR.

| **Primer Name** | **Sequence** | **Gene Target** |
| --- | --- | --- |
| *Bacterial gene expression* | | |
| dnaQ_F | GTCAGGCCCGCAAATTAC | *dnaQ*, endogenous control |
| dnaQ_R | TCTCCACCAGATCGAGACG | *dnaQ*, endogenous control |
| regA_F | TGCGGGATGCTCAACTTCTT | *regA* |
| regA_R | TTTTACGCCGCGTGTGATTG | *regA* |
| ler_F | CGGAGCTCATCAAAAGGGGT | *ler* |
| ler_R | GTTGTCGACCCACTCCAGAC | *ler* |
| eae_F | GGTTAATCTGCAGAGCGGTAA | *eae* |
| eae_R | GAACGGTAATAAGAAGTCCAGTGAA | *eae* |
| tir_F | TTGCATCGACCCAATGGTCA | *tir* |
| tir_R | AGTGCTTTGGATACCCTGCC | *tir* |
| nleB1_F | GCTCCTCTCCAGACTGTTTCC | *nleB1* |
| nleB1_R | GCTCTGGCCTTGCTTCAAAC | *nleB1* |
| espA_F | AATCACCGGCGCTTAACTCA | *espA* |
| espA_R | TTCACGCACAAAGCGAACTG | *espA* |
| fimA_F | TATGTCTGCCCTGTCCCTGA | *fimA* |
| fimA_R | TTAAAATGCACTGTGCCGCC | *fmA* |
| kfcC_F | TGCAACCGAGACGGATAAGG | *kfcC* |
| kfcC_R | TTGCCGGGCGTTTTAATTGG | *kfcC* |
| degP_F | CATTAACACCGCGATCCTC | *degP* |
| degP_R | CCATATTGCTCGGGATAGCA | *degP* |
| cpxA_F | GGTGTTGATGTTGCCCAAGC | *cpxA* |
| cpxA_R | GCCTCGACGTGTTGCTCTAT | *cpxA* |
| rpoS_F | GAAGACACCACGCAGGATGA | *rpoS* |
| rpoS_R | CCGCTTCATAACCCAGCAGA | *rpoS* |
| recA_F | TGCCACTACCTGGCTGAAAG | *recA* |
| recA_R | CGTGGAGTCCTGGTTGTTGA | *recA* |
| *Host gene expression* | | |
| b2m_F | CCAAGACCGTCTACTGGGGT | *B2M*, endogenous control |
| b2m_R | TCTGCTGGCACCACAGATCA | *B2M*, endogenous control |
| tjp1_F | CCCTGAAAGAAGCGATTCAG | *TJP1* |
| tjp1_R | CCCGCCTTCTGTATCTGTGT | *TJP1* |
| atp4b_F | GACGTGTATGGGGAAAGAGGG | *ATP4B* |
| atp4b_R | GATGCTGGGGTATAGCCTGCTAAG | *ATP4B* |
| hrh2_1_F | AAGCCCAGTCAGGACACGAC | *HRH2* |
| hrh2_1_R | TCCCAGCTTCCAGATGAAGGA | *HRH2* |
